# Cholinergic modulation of distinct inhibitory domains in granule cells of the olfactory bulb

**DOI:** 10.1101/2021.10.15.464603

**Authors:** Pablo S. Villar, Ruilong Hu, Batya Teitz, Ricardo C. Araneda

**Author notes:** Corresponding author: Ricardo C. Araneda.

## Abstract

Early olfactory processing relies on a large population of inhibitory neurons in the olfactory bulb (OB), the granule cells (GCs). GCs inhibit the OB output neurons, the mitral and tufted cells (M/TCs), shaping their responses to odors both in the spatial and temporal domains, therefore, the activity of GCs is finely tuned by local and centrifugal excitatory and inhibitory inputs. While the circuit substrates underlying regulatory inputs onto GCs are well-established, how they are locally modulated remains unclear. Here, we examine the regulation of GABAergic inhibition onto GCs by acetylcholine, a main neuromodulatory transmitter released in the OB, by basal forebrain (BF) neurons. In acute brain slices from male and female mice, we show that activation of muscarinic acetylcholine receptors (mAChRs) produces opposing effects on local and centrifugal inhibition onto GCs. By using electrophysiology, laser uncaging and optogenetics we show that the kinetics of GABAergic currents in GCs could be correlated with distal and proximal spatial domains from where they originate, along the GC somatodendritic axis. Proximal inhibition from BF afferents, is suppressed by activation of M2/M4-mAChRs. In contrast, distal local inhibition from deep short axon cells (dSACs) is enhanced by activation of M3-mAChRs. Furthermore, we show that the cholinergic enhancement of distal inhibition in GCs reduces the extent of dendrodendritic inhibition in MCs. Interestingly, the excitatory cortical feedback, which also targets the proximal region of GCs, was not modulated by acetylcholine, suggesting that muscarinic activation shifts the synaptic balance towards excitation in GCs. Together, these results suggest that BF cholinergic inputs to the OB fine tune GC-mediated inhibition of M/TCs by differentially modulating the proximal and distal domains of inhibition in GCs.

## INTRODUCTION

GABAergic inhibition has a crucial role in shaping sensory processing, precisely regulating both temporal and spatial aspects of signal processing in sensory circuits (Isaacson and Scanziani, 2011; Wood et al., 2017; Cardin, 2018). Inhibition by GABAergic granule cells (GCs) is a prominent physiological mechanism in the olfactory bulb (OB), the first brain region where odor processing occurs (Price and Powell, 1970a; Shepherd, 2004; Abraham et al., 2010; Li et al., 2018). GCs synapse onto mitral and tufted cells (M/TCs), the output neurons of the OB, and the GC function is finely regulated by local and top-down signals (Price and Powell, 1970b; Yokoi et al., 1995; Isaacson and Strowbridge, 1998; Schoppa, 1998; Christie et al., 2001; Shepherd, 2004; Matsutani and Yamamoto, 2008). Among these regulatory signals, both glutamatergic feedback from the olfactory cortices and cholinergic inputs from the BF have been shown to influence odor processing by modulating GCs excitability, in the latter case with a predominant contribution of muscarinic acetylcholine receptors (mAChRs) (Castillo et al., 1999; Ghatpande et al., 2006; Pressler et al., 2007; Smith et al., 2015). Furthermore, the excitatory feedback to GCs is regulated by local GABA release (Mazo et al., 2016); however, less is known about the regulation of GABAergic inhibition of GCs.

GCs receive inhibitory inputs from local deep short-axon cells (dSACs) (Pressler and Strowbridge, 2006; Eyre et al., 2008; Burton and Urban, 2015) and from long-range GABAergic neurons (LRGNs) of the BF (Gracia-Llanes et al., 2010; Nunez-Parra et al., 2013; Sanz Diez et al., 2019; Böhm et al., 2020; Hanson et al., 2020; Villar et al., 2021). Interestingly, these two sources of inhibition appear to be segregated along the proximal-distal axis of GCs, suggesting they could differentially regulate GC function. Inputs from dSACs provide distal, and to a lesser extent proximal, feedforward inhibition to GCs (Eyre et al., 2008; Burton and Urban, 2015), while BF-LRGNs densely innervate deeper layers of the OB, where the somas of GCs are localized, suggesting a predominant perisomatic influence in GCs (Gracia-Llanes et al., 2010; Villar et al., 2021). Furthermore, through dendrodendritic synapses with the M/TCs at their distal dendrites, GCs engage in two types of inhibitory mechanisms to regulate M/TCs activity: localized recurrent inhibition (Isaacson and Strowbridge, 1998; Mori et al., 1999; Egger et al., 2003; Shepherd, 2004), and lateral inhibition, which involves widespread activation of the distal dendrite and the inhibition of several neighboring M/TCs. Thus, proximal vs. distal regulation of GC excitability is expected to have a different influence in tunning recurrent and lateral inhibition, with global excitation of GC dendrites having a larger impact on overall inhibition on M/TCs.

Here, we examined the regulation of inhibition to GCs by acetylcholine (ACh). By using electrophysiology, laser uncaging and optogenetics we show that proximal and distal anatomical domains of inhibition can be distinguished in GCs, based on the waveforms of inhibitory currents onto GCs. Events originating in distal regions of the GC’s dendrites corresponded to inputs from dSACs. Accordingly, depolarization of dSACs by activation of M3-mAChRs produced a robust increase in the occurrence of distal sIPSCs. Importantly, the increase in distal inhibition onto GCs decreased recurrent inhibition in mitral cells (MCs). In contrast, inhibitory inputs from the BF, targeting the proximity of the soma, were strongly suppressed by activation of M2-mAChRs. Interestingly, the excitatory feedback from the piriform cortex (PC), which similar to LRGNs targets the somatic region of GCs, was not affected by ACh, suggesting that at the somatic level, ACh promotes excitation of GCs. Together, these results suggest that topologically distinct sources of inhibition of GCs are differentially modulated by ACh through the activation of muscarinic receptors, which we propose will facilitate global excitation of GC, while enhancing inhibition onto distal dendritic compartments.

## RESULTS

### Muscarinic ACh receptor activation increases inhibitory activity in GCs

The BF GABAergic and cholinergic projections to the OB exhibit a considerable overlap in most layers, especially in the GC and periglomerular layers where abundant synaptic boutons can be observed (**Figure 1A, insets**) (mean normalized pixel intensity ± SD, Gad2 MCPO axons: GL 0.3 ± 0.1, EPL 0.3 ± 0.1, GCL 0.6 ± 0.1, n= 14 slices, 3 mice; ChAT BF axons: GL 0.7 ± 0.2, EPL 0.5 ± 0.1, GCL 0.5 ± 0.1, n= 22 slices, 4 mice). The extent of overlap of these systems suggests that as in other brain regions, ACh can modulate GABAergic inhibition in GCs. To examine this possibility, we bath perfused ACh (100 μM) while recording sIPSCs in GCs (**Figure 1B)**. We found that in 50% of the GCs recorded (9/18), ACh strongly increased the frequency of inhibitory events (control 1.1 ± 0.7 Hz vs. ACh 2.4 ± 1.1 Hz, n= 9, p= 0.04) (**Figure 1C**), especially sIPSCs of smaller amplitude (**Figure 1B**, arrowheads). The effect of ACh on GCs was completely abolished in the presence of atropine (Atrp, 3 μM), a broad muscarinic receptor blocker (control 1.2 ± 0.6 vs. ACh in Atrp 1.2 ± 0.5 Hz, n= 10, p= 0.82) (**Figure 1B, C**), indicating the enhancement of inhibition in GCs results from mAChR activation. Accordingly, the mAChR agonist muscarine (Mus, 10 μM), produced a significant increase in the frequency of sIPSCs in ∼50% of the recorded GCs (13/24) (control 0.3 ± 0.08 Hz vs. Mus 1.7 ± 0.55 Hz, n= 13, p= 0.02) (**Figure 1C**). As with ACh, the increase in event frequency produced by Mus was accompanied by a small reduction in the mean amplitude of the overall population of sIPSCs (control 19.6 ± 0.4 pA vs. Mus 17.3 ± 0.3 pA, n= 13 cells, p< 0.001) (**Figure 2-1A**).

**Figure 1.**
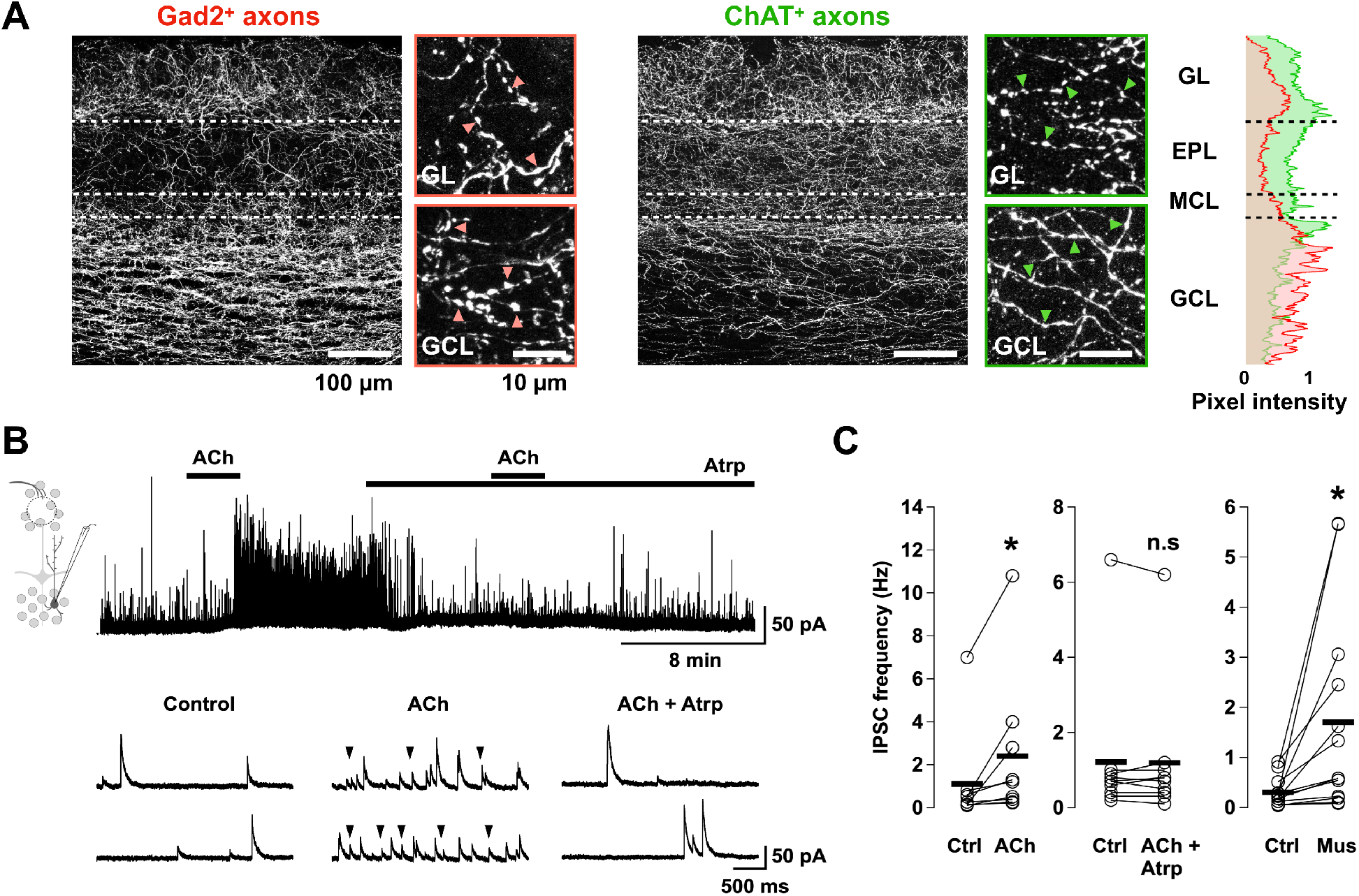
Acetylcholine enhances inhibitory currents in granule cells via muscarinic activation **(A)** Left, horizontal section of the OB of a *Gad2-cre* mouse containing axons from MCPO LRGN expressing eGFP. Right, horizontal section of the OB of a *ChAT-tau GFP* mouse. The panels on the right of each image are magnifications of the GL and GCL showing the presence of abundant axonal boutons (arrowheads) in the GABAergic and cholinergic axons, across the superficial and deeper layers of the OB. Right, quantification of peak normalized pixel intensity distributions for the GABAergic (red) and cholinergic (green) axonal projections shown on the left. There is an extensive colocalization of cholinergic and GABAergic axons, especially in the superficial and deeper layers of the OB. **(B)** Top, spontaneous IPSCs (sIPSCs) were recorded at 0 mV in GCs. In this cell, the basal rate of inhibitory currents was 0.48 Hz. Application of acetylcholine (ACh, 100 μM) produced a large increase in the frequency of sIPSCs (4 Hz), which was prevented by the application of the muscarinic receptor blocker atropine (Atrp, 3 μM, 0.46 Hz). Bottom, selected traces in an expanded time scale, before and after the drugs application shown in the top trace. Arrowheads indicate the increase in frequency of small amplitude events. **(C)** Summary of the effect of ACh and muscarine (Mus, 10 μM) on the sIPSC frequency in GCs; ACh significantly increased the frequency of sIPSC in GCs (n= 9, p= 0.04), and this effect was blocked by Atrp (n= 10, p= 0.82). Similarly, Mus significantly increased the sIPSC frequency in GCs (n= 13, p= 0.02).

In the non-responding GCs, neither ACh nor Mus produced a change in the frequency or the amplitude of the sIPSCs (data not shown). The lack of muscarinic effect in these GCs is unknown, however, both responding and non-responding GCs had a similar basal frequency of sIPSCs (ACh responding 1 ± 0.7 Hz vs. ACh non-responding 0.5 ± 0.1 Hz, p= 0.45; Mus responding 0.3 ± 0.1 Hz vs. Mus non-responding 0.5 ± 0.2 Hz, p= 0.3), arguing against the possibility the non-responding neurons correspond to unhealthy GCs.

### Muscarinic ACh receptor activation enhances distal inhibition of GCs

In the presence of mAChR activation small amplitude events were more prominent, therefore, we examined the possibility that Mus enhanced a selective population of spontaneous GABAergic currents in GCs. Under resting conditions, sIPSCs recorded at 0 mV occurred at low frequency (0.24 ± 0.05 Hz, n= 29) (**Figure 2A**) and exhibited a wide range of amplitudes (5-132 pA, mean ± SD: 23 ± 15 pA, 1411 events), with similar variability in rise (1.1-7.6 ms, mean ± SD: 2.5 ± 2 ms) and decay times (21-145 ms, mean ± SD: 69.1 ± 36 ms), possibly reflecting variations in their origin along the somatodendritic axis (Nusser et al., 1999). To explore this possibility, we clustered the current waveforms of isolated sIPSCs using the k-means method, which rendered three distinct populations of events that we termed clusters 1 to 3 (**Figure 2A** and **Figure 2-1B**). These sIPSC populations had differences in their time course and occurred with different relative frequencies (**Figure 2B, C**). Events in cluster 1, characterized by smaller amplitude, slower rise time and faster decay time, were the most abundant (**Figure 2C**, n= 808 events; mean ± SD: amplitude 13.9 ± 6.3 pA, rise time 3.2 ± 2.7 ms, decay time 62.5 ± 35.4 ms). In contrast, events in cluster 3, characterized by larger amplitude, faster rise times and slower decay, occurred less frequently (**Figure 2C**, n= 150 events; mean ± SD: amplitude 55.2 ± 15.8 pA, rise time 1.3 ± 0.3 ms, decay time 74 ± 29.3). Cluster 2 was characterized by events with intermediate amplitude and occurrence (**Figure 2C**, n= 453 events; mean ± SD: amplitude 28.6 ± 8.3 pA, rise time 1.7 ± 1.4 ms, decay time 79.5 ± 36.3 ms). Interestingly, clustering of the sIPSC waveforms showed that the prevalence of large amplitude sIPSCs (cluster 3) was unaffected by Mus (Wilcoxon rank sum, p= 0.72), while the largest proportional increase occurred with the smaller amplitude sIPSCs (cluster 1 and 2, Wilcoxon rank sum; cluster 1, p= 0.044, cluster 2, p= 0.045) (**Figure 2D**). Consistently, the ratio of sIPSCs after bath perfusion of Mus, compared with baseline, was higher for cluster 1 than for cluster 2 and 3 (**Figure 2E**).

**Figure 2.**
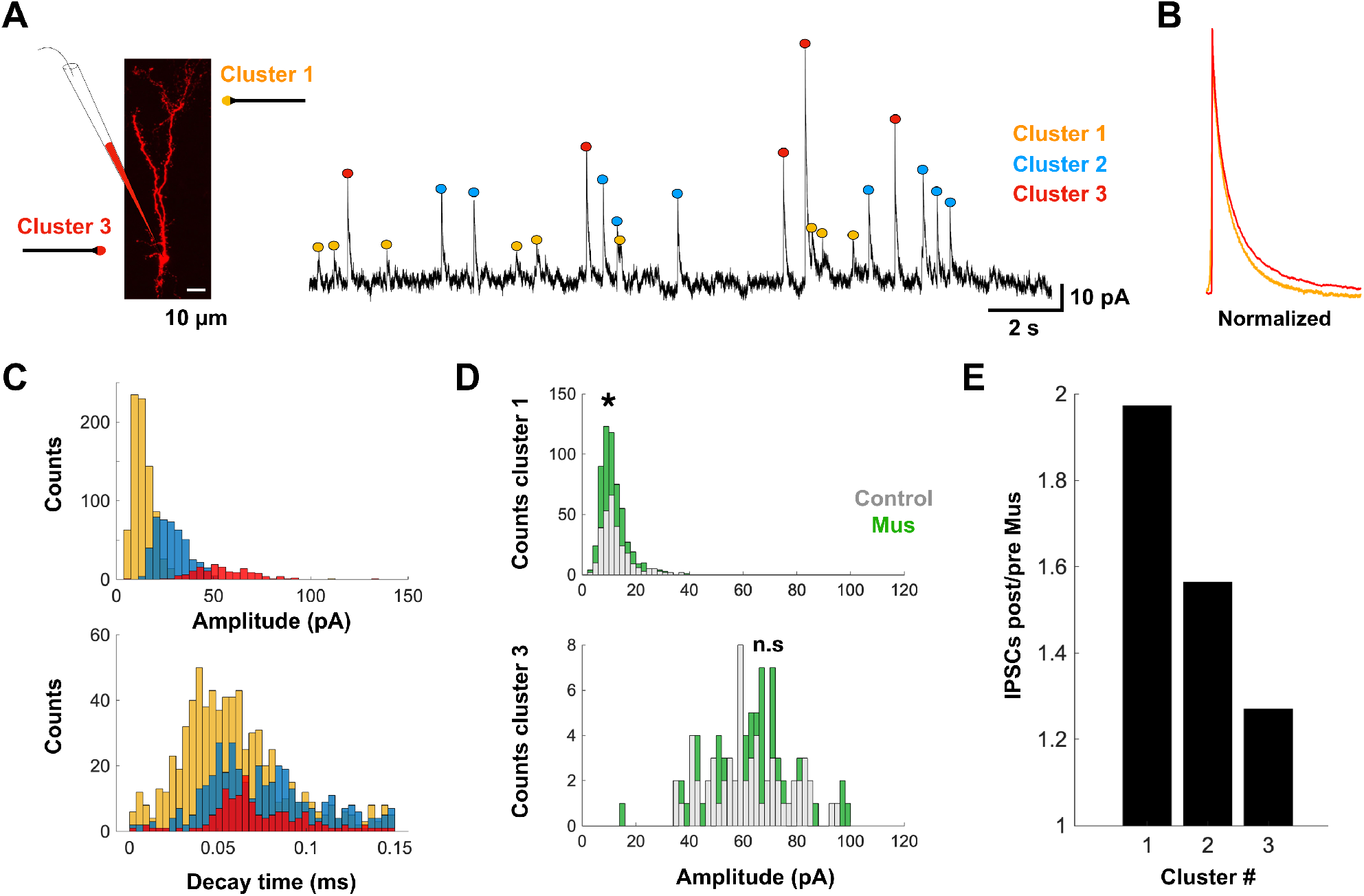
Differential enhancement of inhibitory currents in granule cells by muscarinic AChR activation **(A)** Left, confocal image of a GC filled with Alexa fluor-594 during the whole-cell recording highlighting the presence of intact dendritic processes. Right, sample trace of spontaneous inhibitory postsynaptic currents (sIPSCs) recorded in a representative GC. Recordings were conducted in voltage clamp at 0 mV, using a Cs-gluconate based internal solution. sIPSC waveforms were clustered into three populations using the k-means method. The colored markers on top of each sIPSC indicate the assigned cluster (yellow, cluster 1; blue, cluster 2; red, cluster 3). **(B)** Overlay of normalized waveforms (to the peak current) for clusters 1 and 3 which exhibit different rise and time kinetics. **(C)** Histograms of sIPSC amplitude (top) and decay time (bottom) for each cluster. **(D)** Effect of Mus on events classified as belonging to clusters 1 and 3 (1420 events); the sIPSC amplitude histograms for each cluster are shown, before (light gray) and after the application of Mus (green). Mus significantly increased the number of sIPSC in cluster 1 (p= 0.044) but not in cluster 3 (p= 0.72). **(E)** Summary plot showing the ratio for the occurrence of IPSCs during the application of Mus (post) compared with baseline (pre) (post/pre ratio, cluster 1: 1.97, cluster 2: 1.56, cluster 3: 1.27). The extended **Figure 2-1** shows the un-clustered amplitude distribution for sIPSCs in GCs and mean waveforms for each cluster.

Oxotremorine (Oxo, 10 μM), another non-selective mAChR agonist, produced a similar increase in sIPSC frequency in 50% of GCs (events in clusters 1 and 2; control 0.7 ± 0.3 Hz vs. Oxo 1.4 ± 0.4 Hz, p= 0.05, n= 4 out of 8 cells; not shown), without altering the representation for events in cluster 3. In summary, the clustering method described above allows us to distinguish at least two distinct populations of sIPSCs, clusters 1 and 3. We surmise that cluster 1, with smaller amplitude events, corresponds to sIPSCs originating at distal regions of GC, while events comprising cluster 3, with larger amplitudes, originate at proximal sites; activation of mAChRs mostly increases inhibitory currents originating at distal regions of GCs (see also below). Events in cluster 2, likely correspond to an overlap of inputs that mainly target these proximal and distal regions of GCs; therefore, we concentrated our analysis on clusters 1 and 3.

### Inputs to GCs from dSACs and BF-LRGNs generate IPSCs with distinct properties

GCs receive inhibitory inputs from dSACs found in the external plexiform layer (EPL-dSAC) that directly synapse onto the distal dendrites of GCs (Eyre et al., 2008), and from LRGNs of the BF, which densely innervate deeper layers in the bulb suggesting a greater influence on the somatic region of GCs (Villar et al., 2021). We reasoned that events in clusters 1 and 3 correspond to these sources of inhibition to GCs and examined this possibility by selectively activating these inhibitory inputs using optogenetics. We activated dSACs by stimulating M/TCs expressing the light-gated cation channel channelrhodopsin-2 (ChR2), under the *Thy1* promoter (Arenkiel et al., 2007) (**Figure 3A, B**). To isolate the inhibitory input from dSACs we recorded GCs at the reversal potential of glutamatergic currents (0 mV). Consistent with their disynaptic nature, brief light pulses (0.5-1 ms), reliably evoked IPSCs with slow onset (9 ± 0.2 ms). The evoked IPSCs (n= 451 events, n= 8 cells) had variable amplitude (5-108 pA, mean 23 ± 0.8 pA) and a mean decay of 38.9 ± 1.3 ms (**Figure 3E**).

**Figure 3.**
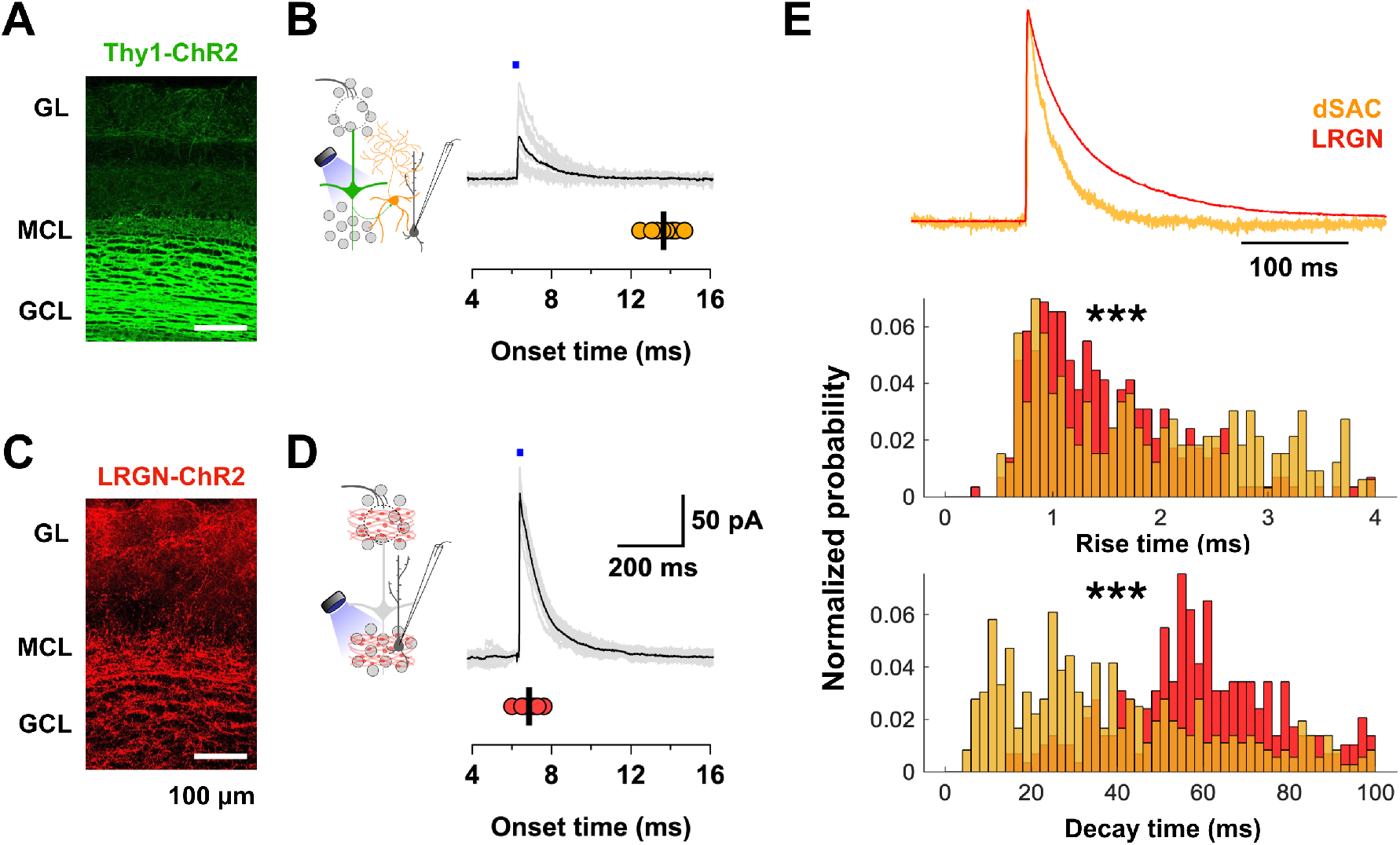
Centrifugal and local GABAergic inputs on granule cells exhibit distinct kinetics **(A)** Confocal image of an OB section showing the expression of ChR2-YFP in Thy1^+^ neurons (M/TCs). **(B)** Left, diagram of the experimental configuration to elicit disynaptic inhibition in GCs; we optogenetically stimulated M/TCs in a *Thy1-EYF-ChR2* mouse while recording at 0 mV. Right trace, overlay of light-evoked disynaptic IPSCs in a representative GC (gray traces) elicited by feedforward excitation of dSACs, the mean evoked IPSC is shown overlaid in black. Bottom, scatter plot of the onset time for the responses in this GC; the average onset time was 13.7 ± 0.2 ms, consistent with a disynaptic nature. **(C)** Confocal image of an OB section showing abundant axonal labeling from MCPO LRGNs expressing ChR2-tdTomato, in a *Gad2-Cre* mouse. **(D)** Left, diagram of the experimental configuration; we recorded from GCs at 0 mV, eliciting minimal optogenetic stimulation of LRGN axons expressing ChR2 (in red). Right trace, overlay of light-evoked IPSCs in a representative GC (gray traces), the mean evoked IPSC is shown overlaid in black. Bottom, scatter plot of the onset times for the responses in the cell above; the average onset time was 7 ± 0.8 ms, consistent with their monosynaptic nature. **(E)** Top, overlay of normalized waveforms (to the peak current) for the dSAC evoked IPSC (yellow) and LRGN evoked IPSC (red) recorded in GCs to illustrate their different decay kinetic. Bottom, probability distribution histograms of rise and decay times for minimally evoked IPSCs with stimulation of BF-LRGNs axons (red; events= 291; n= 10 cells) and the disynaptically evoked IPSC (yellow, events= 451; n= 8 cells). The disynaptic IPSC elicited by dSAC activation exhibits faster decay times, as well as a slightly slower rise times compared with the LRGN evoked IPSC (dSAC evoked IPSC: decay time 38.89 ± 1.3 ms, rise time 1.9 ± 0.05 m; MCPO evoked IPSC: decay time 59.1 ± 1 ms, rise time 1.5 ± 0.04 ms. Wilcoxon rank sum p< 0.001).

To selectively activate the BF GABAergic inputs to GCs, we expressed ChR2 in the MCPO of *Gad2-Cre* mice (Villar et al., 2021). Light activation evoked IPSCs (n= 291 events, 10 cells) of faster onset (7 ± 0.8 ms), larger amplitude (mean 130 ± 5.5 pA), and slower decay (mean 59.1 ± 1 ms), compared to those evoked by dSACs activation. As shown in Figure 3E, the normalized currents for the dSAC mediated inhibitory events and those elicited by LRGN activation, show a significant difference in rise and decay times, reminiscent of the events in cluster 1 and 3 respectively (comparison for amplitude, rise time and decay time Wilcoxon rank sum p< 0.001). Together, these results suggest that sIPSCs in cluster 1 correspond to events elicited by GABA release from dSACs, preferentially targeting the distal dendrites of GCs, whereas events in cluster 3 represents inhibitory inputs from BF-LRGN targeting the perisomatic region of GCs.

To further corroborate that the differences in amplitude and kinetics of the somatically recorded sIPSCs correspond to their spatial origin, we analyzed the properties of GABA evoked currents in GCs using single-photon uncaging (**Figure 4A**). In these experiments, the photolysis of caged GABA (DPNI-GABA) was calibrated to obtain photo-evoked inhibitory postsynaptic currents (uIPSC) of similar amplitude to the sIPSCs (∼10-100 pA). To visualize the dendritic arbor of GCs, we included the red fluorophore Alexa fluor-594 (20 μM) in the internal solution, and the stimulation was ∼10 μm apart along the GC somatodendritic axis (**Figure 4A**; see Methods). GABA uncaging in the perisomatic region (0-20 μm), resulted in uIPSCs with an average amplitude of 79.1 ± 10.6 pA and a decay time of 360 ± 30 ms, while at most distal regions (150-250 μm from the soma) both the amplitudes and decay times of the uIPSC were smaller (14.1 ± 1.4 pA; 170 ± 50 ms; n= 4 cells) (**Figure 4B, C**). In contrast, the rise time was not significantly different with distance (proximal 25 ± 4 ms vs distal 36 ± 10 ms, n= 4 cells, p= 0.24) (**Figure 4C**). Together, these results further support the notion that sIPSCs in cluster 1, enhanced in frequency by Mus, correspond to events elicited by GABA release from dSACs and preferentially target the distal dendrites of GCs, whereas events in cluster 3 represents inhibitory inputs from BF-LRGN targeting the perisomatic region of GCs.

**Figure 4.**
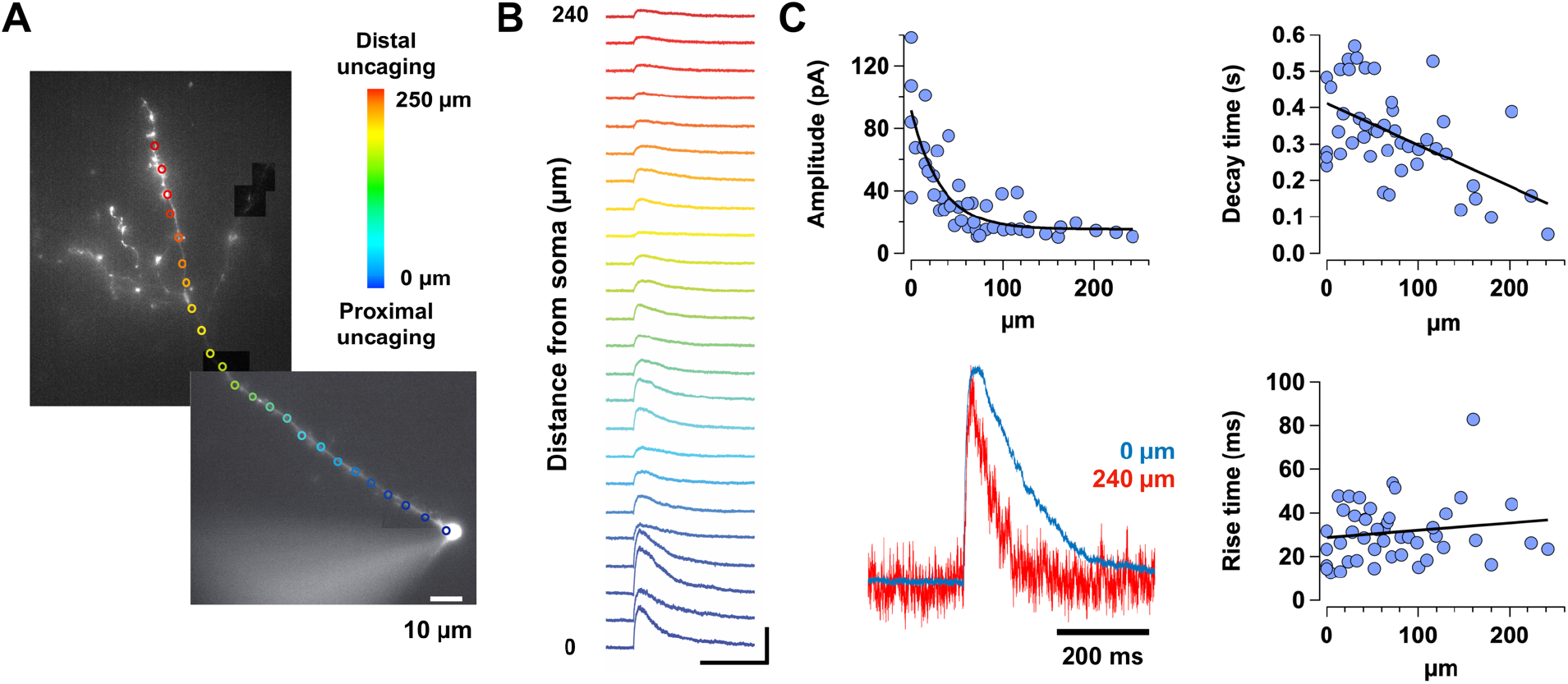
Spatial differences in the kinetics of GABAergic currents elicited by laser uncaging along the GC somatodendritic axis **(A)** Representative GC loaded with Alexa fluor-594 and recorded in voltage-clamp, at a holding potential of 0 mV. DPNI-GABA was uncaged along the somatodendritic axis (colored circles) with a 405 nm laser focused through a 60x objective. Cold colors represent proximal spots while warm colors represent distal focused spots. **(B)** Mean traces for the IPSCs elicited by GABA uncaging at different distances from the soma. The color of each trace corresponds to the uncaging position as shown in A. The scale is 100 pA and 200 ms. **(C)** Scatter plots of the amplitude, and the rise and decay time as a function of distance from the soma (n= 4 cells). The amplitude and the decay time of the responses (top graphs) decreased as GABA was uncaged further away from the soma (amplitude: double exponential fit, R-squared= 0.675; decay time: Pearson’s linear correlation, r= –0.55). Lower left graph, overlay of representative normalized (to the peak current) traces obtained by uncaging in the proximal (blue trace) and distal (red trace) regions of a GC dendrite illustrating their different kinetics. In contrast, the rise time was not affected by the uncaging distance (bottom right, Pearson’s linear correlation, r= 0.15).

### dSAC are excited by M3-mAChR activation

To further examine the possibility that the increase in distal inhibition results from dSACs activation by Mus, we recorded from morphologically and physiologically identified dSACs loaded with Alexa fluor-594 in the recording pipette (**Figure 5A**). Morphological analysis indicated that these cells exhibit numerous basal dendrites as well as axonal projections directed toward the EPL (see also Eyre et al., 2008). As previously described (Pressler and Strowbridge, 2006), under resting conditions, a current-stimulus elicited a train of spikes that was followed by an afterdepolarization (ΔV 2 ± 0.8 mV; n= 4), (**Figure 5A, arrow**). As shown in **Figure 5B**, application of Mus produced a robust depolarization of dSACs (ΔV= 5.7 ± 1.1 mV, n= 4, p= 0.01), which was abolished by application of the selective M3-mAChR antagonist 4-DAMP (100 nM, **Figure 5C**, Mus ΔV 5.74 ± 3.7 mV, Mus in 4-DAMP ΔV 0.35 ± –0.52 mV, n= 4, p= 0.04). Importantly, the disynaptically evoked IPSC recorded in GCs upon M/TCs activation using the *Thy1-EYF-ChR2* mouse was not significantly affected by the activation of mAChRs (control 5.2 ± 1.5 pC vs. Mus 3.9 ± 1 pC, n= 7, p= 0.3) (**Figure 5D, E**).

**Figure 5.**
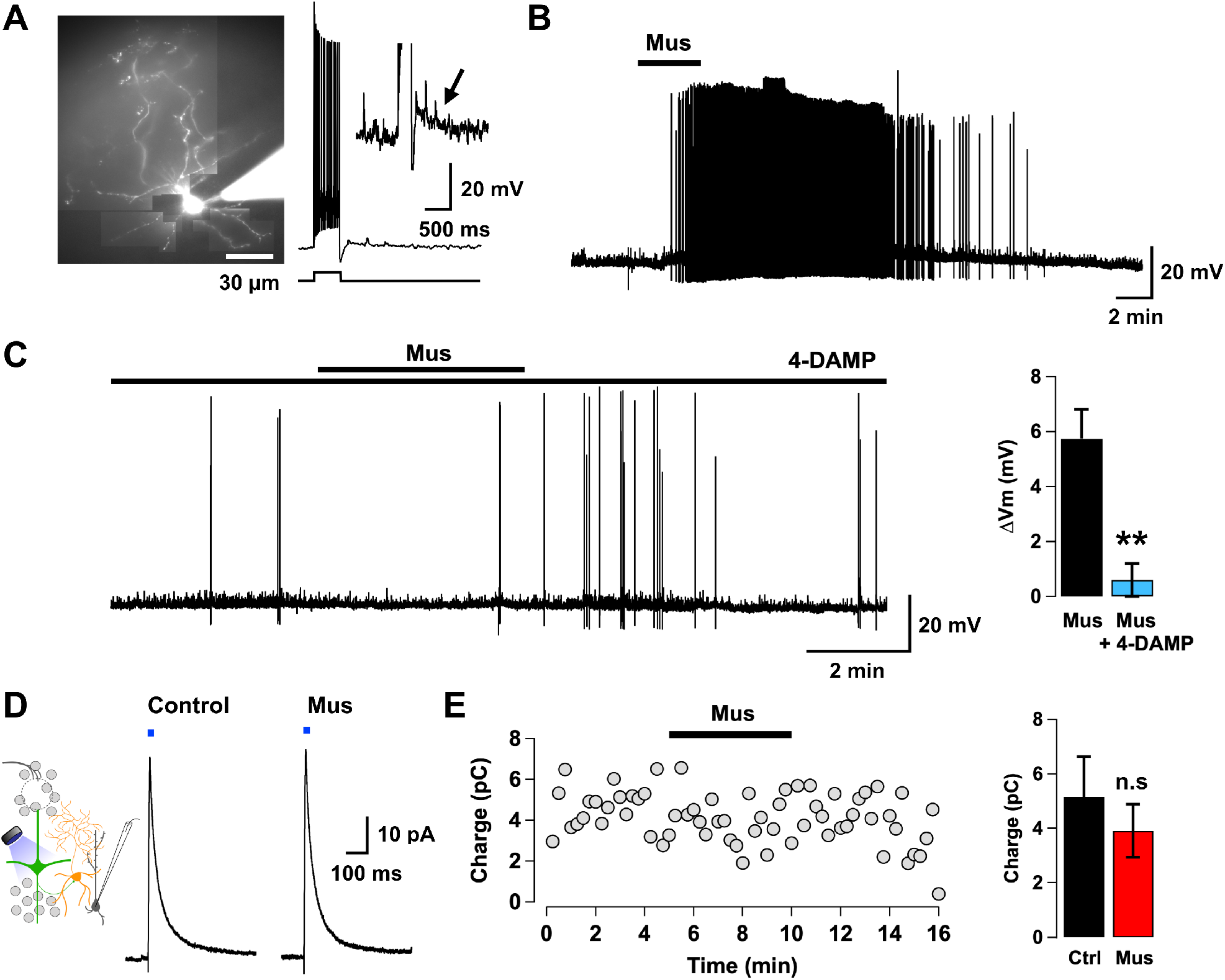
Deep short axon cells are excited by M3-mAChR activation **(A)** Left, image of a dSAC loaded with Alexa fluor-594 during a whole-cell recording. Right, these cells exhibit an afterdepolarization (inset, arrow) following a stimulus-induced train of action potentials (150 pA, 500 ms). **(B)** Application of muscarine (Mus, 10 μM) produced a suprathreshold depolarization in this dSACs, with long-lasting spiking. The baseline membrane potential is –60 mV. **(C)** Left, the muscarinic depolarization was significantly reduced in the presence of 4-DAMP (100 nM), a selective M3-mAChR blocker. Right, summary of the effects of Mus and the M3 antagonist in dSACs (n= 4, p= 0.01). (**D**) Diagram showing the experimental configuration used to examine the effect of Mus on the dSAC-mediated inhibition of GCs. Optogenetic stimulation of M/TCs in the *Thy1-EYF-ChR2* mouse reliably evoked disynaptic IPSCs in GCs recorded at 0 mV. (**E**) The charge of the IPSC evoked by optogenetic stimulation of M/TCs, is not significantly different in the presence of Mus (n= 7, p= 0.3).

Depolarization of dSACs by M3-mAChRs activation, is expected to increase GABA release onto GCs, and accordingly, application of 4-DAMP (100 nM) completely abolished the increase in sIPSC frequency in GCs elicited by Mus (**Figure 6A, B**, control 0.54 ± 0.19 Hz vs. Mus in 4DAMP 0.49 ± 0.14 Hz, n= 11, p= 0.41). As expected, the enhancement of inhibition in GCs produced by Mus was not prevented by selective blockade of M1- and M2/M4-mAChRs with pirenzepine (Pir, 1 μM) and AFDX-384 (300 nM), respectively (**Figure 6B**, control in Pir + AFDX-384 0.46 ± 0.1 Hz vs. Mus in Pir + AFDX-384 0.79 ± 0.2 Hz, n= 6, p= 0.04). Together, these results indicate that distal inhibition is enhanced in GCs via an excitatory effect on dSACs that is mediated by M3-mAChRs activation.

**Figure 6.**
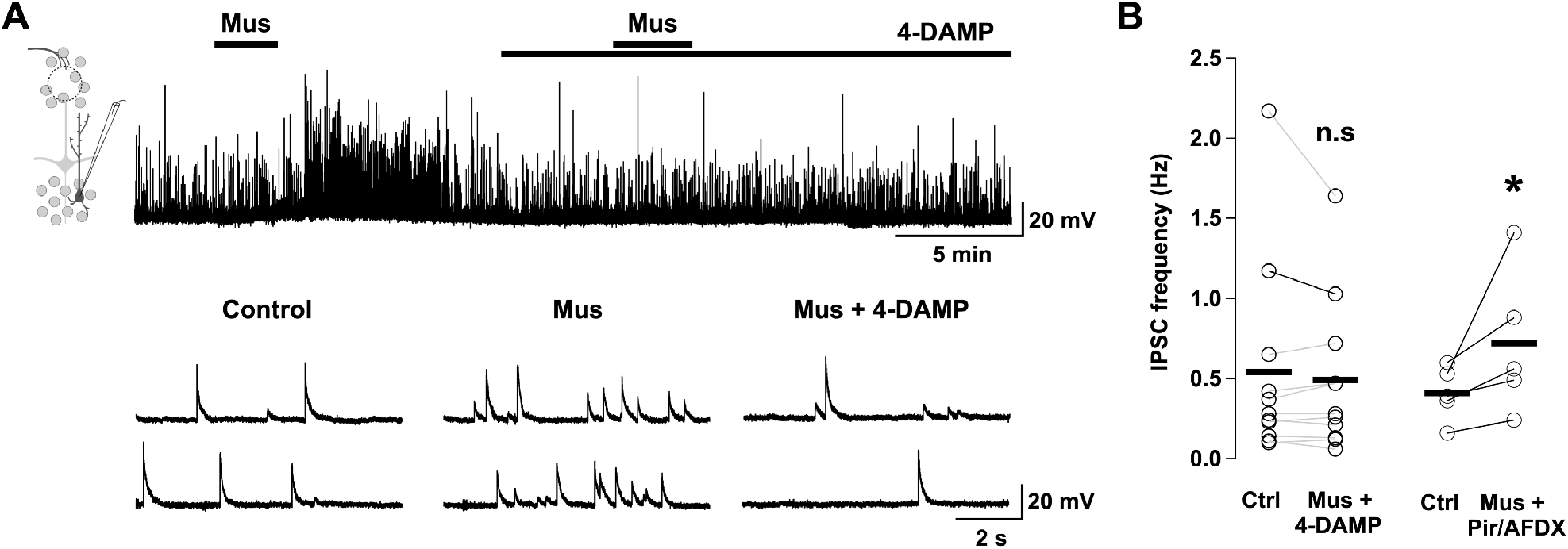
M3-mAChR activation is necessary for the enhancement of inhibition in GCs **(A)** Top, the increase in sIPSC frequency in GCs produced by muscarine (Mus, 10 μM) is abolished in the presence of the selective M3-mAChR blocker 4-DAMP (100 nM). The holding potential is 0 mV. Bottom, selected traces of sIPSCs in an expanded time scale, before and after the drug applications as shown in the top trace. **(B)** Summary plots showing the effect Mus and specific mAChR antagonists. Left, 4-DAMP abolishes the effect of Mus (n= 11, p= 0.4); however, the effect of Mus is not antagonized by a mixture of the M1 and M2/4-mAChR blockers pirenzepine (Pir, 1 μM) and AFDX-384 (300 nM), respectively (n= 6, p= 0.04).

### Activation of M2/M4-mAChRs suppresses proximal inhibition onto GCs

We next examined the effect of ACh (100 μM) on inhibitory currents elicited in GCs by optogenetic activation of BF-LRGNs axons in the OB. In contrast to the effect on distal inhibition, ACh produced a significant reduction in the light-evoked IPSCs (**Figure 7A**, control 14 ± 4 pC vs. ACh 2.8 ± 0.9 pC, n= 7, p= 0.03), and this effect was completely blocked by Atrp (**Figure 7A, B** control 11 ± 2.2 pC vs. ACh in Atrp 11 ± 2 pC, n= 6, p= 0.8). Likewise, Mus (10 μM) produced a strong reduction in the light-evoked IPSCs (**Figure 7C**, control 9.0 ± 2.3 pC vs. Mus 1.5 ± 0.4 pC, n= 4, p= 0.05), and this effect was blocked by AFDX-384 (300 nM), a selective M2/M4-mAChR antagonist (**Figure 7C**, control, 7.5 ± 1 pC vs. Mus + AFDX-384, 6 ± 0.6 pC, n= 4, p= 0.1), suggesting a presynaptic inhibitory action of M2/M4-mAChRs activation on GABA release from BF-LRGNs axons.

**Figure 7.**
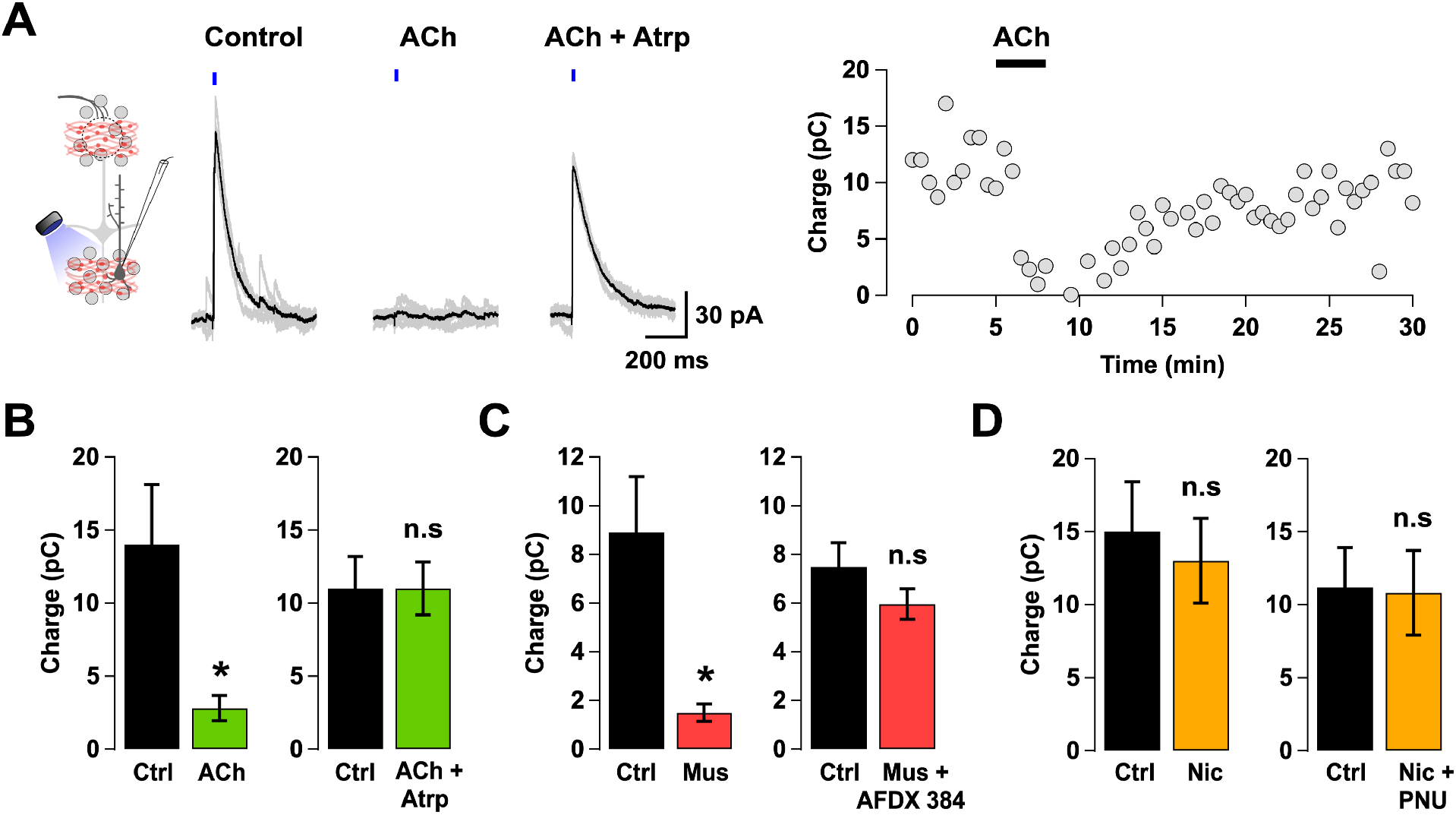
Activation of M2/M4-mAChR suppresses afferent inhibition onto GCs **(A)** Recordings were conducted in GCs at 0 mV, in slices of mice expressing ChR2 in MCPO LRGNs (shown in red in the diagram). Left, stimulation with a brief light pulse (1-5 ms) reliably elicited IPSCs in GCs (gray traces); the average amplitude is 122 pA in this cell (black trace). In the presence of ACh (100 μM), optogenetic stimulation failed to evoke the IPSC. The suppressive effect of ACh was blocked by the muscarinic antagonist atropine (Atrp, 3 μM). Right, time course of the effect of ACh in a representative GC where light-evoked IPSCs occurred every 30 s; ACh reversibly decreased the charge of the IPSC. **(B)** Summary of the effects of ACh on the light-evoked IPSC (left, n= 7, p= 0.03) and the blockade by Atrp (right, n= 6, p= 0.8). **(C)** Left, summary of the effect of Mus (10 μM) on the light-evoked IPSC (n= 4, p= 0.05). Right, the suppression of the evoked IPSC by Mus is reversed in the presence of the M2/M4-mAChR antagonist AFDX-384 (300 nM) (n= 4, p= 0.1). **(D)** The light-evoked IPSC was not affected by nicotine (Nic, 60 μM; n= 6, p= 0.11), or by the application of Nic together with the nAChR positive allosteric modulator PNU-120596 (PNU, 10 μM; n= 5, p= 0.74).

In contrast, bath perfusion of the nicotinic ACh receptor (nAChR) agonist nicotine (Nic, 60 μM) had no effect on the light-evoked IPSC in GCs (**Figure 7D**, control 14.5 ± 3 pC vs. Nic 13.4 ± 3 pC, n= 6, p= 0.11). Even in the presence of PNU-120596 (PNU, 10 μM), a positive allosteric modulator of nAChRs that reduces their desensitization (Hurst et al., 2005), Nic did not affect the inhibitory currents elicited in GCs by light stimulation (**Figure 7D**, control 11.2 ± 3 pC vs. Nic + PNU 10.8 ± 3 pC, n= 5, p= 0.74). Together, these results indicate that BF GABAergic inhibition of GCs is negatively modulated by cholinergic inputs to the OB, via activation of mAChRs in the LRGN axon terminals, and that local and top-down inhibition onto GCs are differentially modulated by the cholinergic innervation through the activation of distinct mAChRs.

### M3-mAChR activation reduces the extent of dendrodendritic inhibition in MCs

Increased inhibition at the distal dendritic segments of GCs is expected to negatively impact the extent of inhibition at GCs to MCs synapses. To examine this possibility, we recorded inhibitory currents in MCs evoked by a brief depolarizing train, while holding the cell at −60 mV, using a CsCl based internal solution. This stimulation elicited a barrage of GABAergic currents with a relaxation time of 564 ± 176 ms (n= 9 cells), similar to values previously reported (Isaacson and Strowbridge, 1998; Schoppa, 1998; Villar et al., 2021). In the presence of Mus, dendrodendritic inhibition (DDI) in MCs was significantly reduced (**Figure 8A** control: –55 ± 7 pC vs. Mus: –40 ± 4 pC, n= 18, p= 0.04). Intriguingly, in the presence of Mus, application of the M3-mAChR blocker, 4-DAMP, produced a significant increase in DDI in MCs (**Figure 8B**, control –39 ± 4 pC vs. Mus in 4-DAMP –51 ± 7 pC, n= 16, p= 0.01). We suspect this is due to an increase in excitability in GCs produced by M1-mAChRs activation (Smith et al., 2015), and accordingly, blockade of both M1 and M3-mAChRs abolished this effect (control, –45 ± 8 pC vs. Mus + 4-DAMP + Pir, –42 ± 9 pC, n= 13, p= 0.44; not shown). Last, the reduction in inhibition produced by Mus persisted in the presence of the selective M2/M4 and M1-mAChR blockers, AFDX-384 and Pir, respectively (**Figure 8C**, control in AFDX-384 + Pir: –69 ± 10 pC vs. Mus + AFDX-384 + Pir: –42 ± 9 pC, n= 11, p< 0.001). These results support the hypothesis that the activation of M3-mAChRs in dSAC reduces the extent of DDI in MCs by increasing incoming GABAergic inhibition in the distal dendrites of GCs.

**Figure 8.**
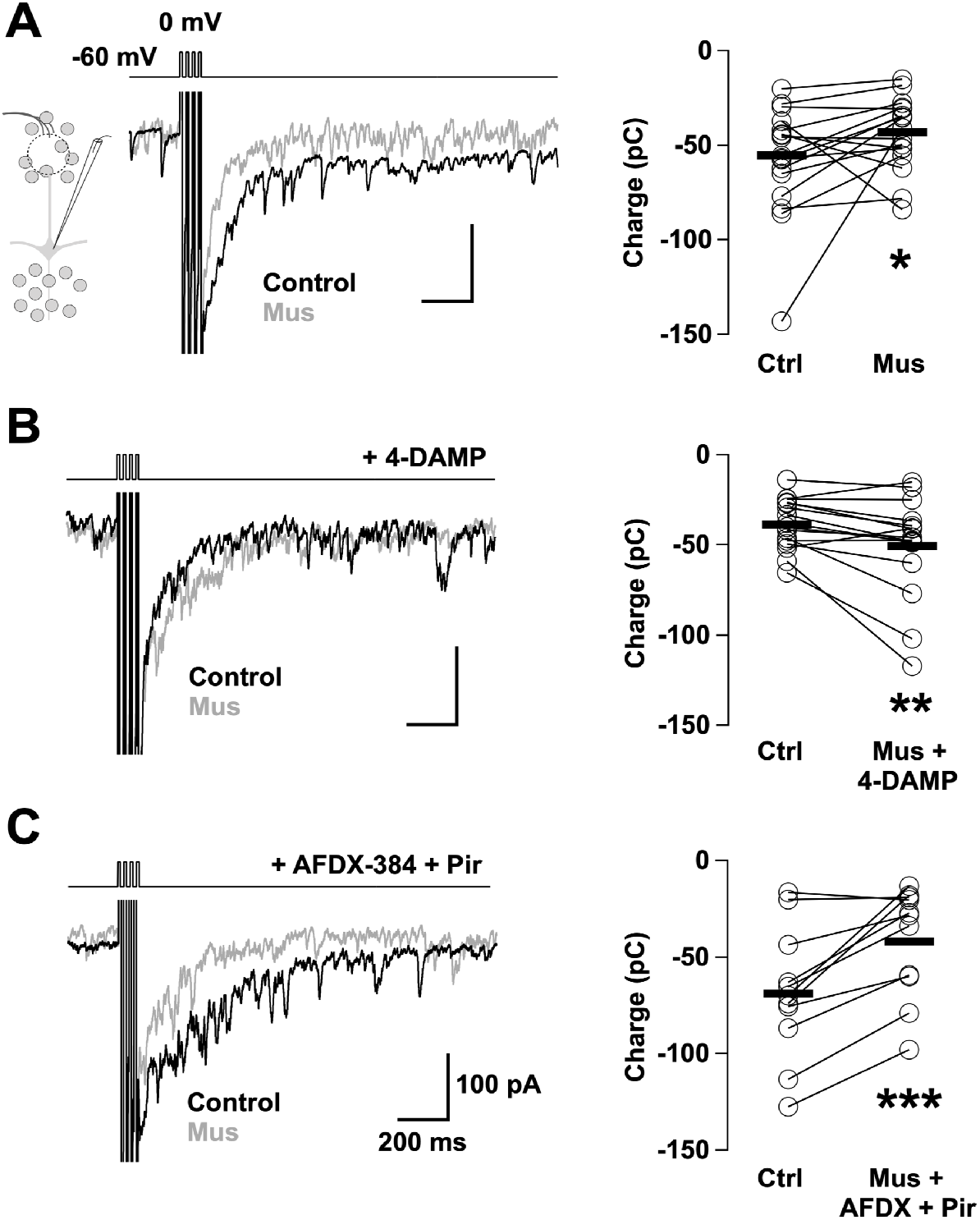
Activation of M3-mAChRs modulates the extent of dendrodendritic inhibition in MCs **(A)** MCs were recorded in voltage-clamp at –60 mV using a CsCl based internal solution; dendrodendritic inhibition (DDI) was elicited by a brief depolarizing train of depolarizing steps (4 x10 ms). The average evoked GABAergic current (black trace) was decreased in the presence of Mus (gray trace, Mus 10 μM). Right, summary of the effect of Mus in DDI quantified as the charge (n= 18, p= 0.04). **(B)** Left, in the presence of the M3-mAChR antagonist, 4-DAMP (100 nM), Mus failed to reduce DDI, but instead produced a small but significant increase in DDI. These effects are summarized in the right plot (n= 16, p= 0.01). **(C)** Left, the decrease in DDI elicited by Mus was still present in the presence of AFDX-384 (300 nM) and pirenzepine (Pir, 1 μM), antagonists of the M2 and M1-mAChRs, respectively. These effects are summarized in the right plot (n= 11, p< 0.001).

### Inhibitory and excitatory top-down inputs onto GCs are differentially modulated by mAChRs

The OB integrates glutamatergic feedback from the PC, which decorrelates the activity of output neurons by driving strong excitation onto the OB local GABAergic circuits (Boyd et al., 2012; Otazu et al., 2015). This excitatory feedback, like the BF inhibitory input, mainly targets the proximal region of GCs (Balu et al., 2007), and therefore, we wondered whether this excitatory input was also inhibited by muscarinic activation. To this extent, we injected the adenovirus AAV-ChR2-mCherry into the PC of wild type mice, which resulted in abundant axonal labelling distributed across the different OB layers, especially in the GCL (**Figure 9A**). In voltage-clamp recordings brief light stimulation (1-5 ms) reliably evoked large inward currents in GCs, which were sensitive to the AMPA receptor blocker CNQX (10 μM) (control –1.3 ± 0.2 pC vs. CNQX –0.2 ± 0.05 pC; n= 7, p= 0.001), as previously shown (Boyd et al., 2012). Intriguingly, application of Mus did not alter the light-evoked EPSC (control –1.3 ± 0.3 pC vs. Mus –1.3 ± 0.2 pC, n= 6, p= 0.88) (**Figure 9B, C**). Similar results were obtained when we applied Nic (60 μM) alone or in the presence of PNU (10 μM) (control, –0.9 ± 0.2 pC vs. Nic, –0.9 ± 0.2 pC, n= 5, p= 0.2; control, –1.1 ± 0.2 pC vs. Nic + PNU, –1 ± 0.3 pC, n= 4, p= 0.6) (**Figure 9C**). Nevertheless, consistent with a previous study (Mazo et al., 2016), activating GABA_B_Rs with the agonist baclofen (Bac, 50 μM) strongly suppressed the light-evoked EPSC in GCs (control –0.9 ± 0.2 pC vs. –0.2 ± 0.1 pC, n= 6, p= 0.02; not shown). These results suggest that the excitatory/inhibitory balance in GCs can be modulated by mAChR activation.

**Figure 9.**
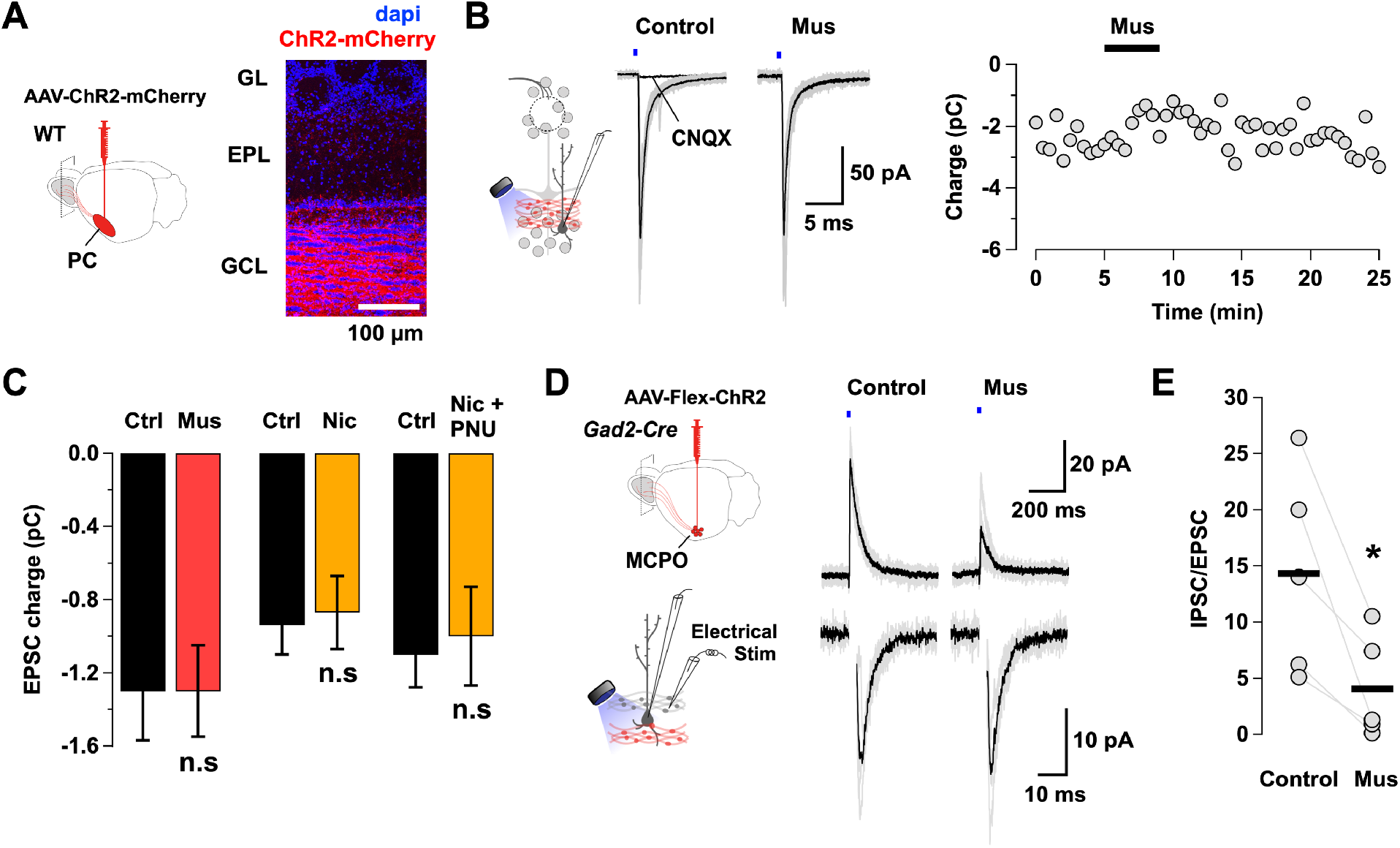
Activation of mAChRs suppresses extrinsic inhibition but not extrinsic excitation of GCs **(A)** Left, cortical feedback projections were transduced with ChR2 by injecting the anterograde virus AAV-ChR2-mCherry in the PC of wild type mice, which resulted in extensive labeling of the GC layer (right). **(B)** We conduced voltage-clamp recordings in GCs while optogenetically stimulating the PC excitatory axons (red). Stimulation with a brief light pulse (1-5 ms) in this GC elicited large inward currents (gray traces), with an average amplitude of 170 pA (black trace). The inward current was completely abolished by the AMPA receptor blocker CNQX (10 μM, n= 7, p= 0.001) but not significantly affected by Mus (10 μM), shown on the right-hand side plot. **(C)** Left, bar graphs summarizing the effects of different cholinergic drugs on the light-evoked EPSC. The PC excitatory input was not affected by Mus (n= 6, p= 0.88), Nic (60 μM, n= 5, p= 0.2) or Nic in the presence of PNU-120596 (PNU, 10 μM) (n= 4, p= 0.6). **(D)** We recorded simultaneously both the afferent excitatory and inhibitory inputs onto GCs in *Gad2-Cre* mice injected with AAV-Flex-ChR-mCherry in the MCPO. The cortical feedback was stimulated by placing an electrode in the GCL (bottom). Right, in this GC, light stimulation at 0 mV elicited outward currents (upper gray traces) with an average of 46 pA (black trace), which was greatly reduced by Mus (18 pA). In the same GC, recorded at –60 mV, brief electrical stimulation in the GCL (100 μA, 100 μs) evoked inward currents (lower gray traces) with an average of 20 pA (black trace), which was not affected by Mus (21 pA). **(E)** The IPSC to EPSC ratio of the synaptic charge (absolute value) evoked by light (inhibition) and electrical stimulation (excitation) in the same GC decreases in the presence of Mus (n= 5, p= 0.02).

Thus, in a subset of GCs, we optogenetically activated the MCPO GABAergic axons (as in Figure 3 and 7), while the cortical feedback was electrically stimulated by placing a stimulating electrode in the GCL (**Figure 9D**) (Balu et al., 2007). The optogenetically evoked IPSC (top trace) and electrically evoked EPSC (bottom trace) were isolated by recording at 0 and –60 mV, respectively, using a Cs-based internal solution. At –60 mV, brief current pulses (100 μA, 100 μs) reliably evoked EPSCs in GCs, which were completely blocked by CNQX (control –0.4 ± 0.2 pC vs. CNQX –0.1 ± 0.05 pC, n= 5, p= 0.05; not shown). In the same GC, Mus selectively suppressed the light-evoked IPSCs, but not the electrically evoked EPSC (IPSC control, 2.6 ± 0.2 pC vs. in Mus, 0.6 ± 0.2 pC n= 5, p< 0.01; EPSC, control, –0.3 ± 0.1 pC vs. in Mus, –0.3 ± 0.1, n= 5, p= 0.7). Consistently, Mus produced a strong decrease in the IPSC to EPSC ratio in GCs (**Figure 9E**) (|IPSC/EPSC|, control 14.3 ± 4 vs. Mus 4 ± 2, n= 5, p= 0.02), suggesting an increase in the overall proximal excitatory weight in GCs upon cholinergic activation.

## DISCUSSION

Here, we provide new mechanistic insights on how synaptic inputs onto proximal and distal dendritic domains of GCs are modulated by the cholinergic system and shape the output of the bulb. Analysis of the waveform of spontaneously occurring GABAergic currents revealed at least two groups of events with distinguishing features; events of small amplitude with fast decay (cluster 1), and events of large amplitude with slower decay (cluster 3). Photo-uncaging of GABA indicated that events in cluster 3 originate in the proximity of the soma, while events in cluster 1 originate in distal regions of GCs. Furthermore, optogenetic activation of dSACs or BF-LRGN axons, elicited IPSCs with properties resembling distal and proximal inputs, respectively, suggesting a spatial segregation of inhibitory inputs in GCs. Interestingly, activation of mAChRs produced opposite effects on these inhibitory inputs; activation of M3-mAChRs greatly enhanced the occurrence of inhibitory currents in GCs, by depolarization of dSACs that synapse onto GCs. In contrast, activation of the M2/M4-mAChRs depressed GABAergic currents elicited by activation of BF inputs. These distinct effects on proximal vs. distal inhibition in GCs are expected to regulate the degree of inhibitory output at M/TC-GC synapses. Activation of M3-mAChRs reduced the extent of dendrodendritic inhibition in MCs, suggesting a local regulation of inhibition. Interestingly, the glutamatergic feedback to the OB was not affected by Mus, suggesting that the overall effect of proximal muscarinic modulation is to shift the balance towards excitation of GC, enhancing global inhibition.

The activity of M/TCs is functionally regulated by several types of inhibitory neurons, which target their primary and secondary dendrites (Price and Powell, 1970a; Shepherd, 2004). Among the inhibitory neurons of the OB, the most prominent in number are the GCs, which participate in recurrent and lateral inhibition through synapses with lateral dendrites of M/TCs (De Olmos et al., 1978; Luskin and Price, 1983). Most dendrodendritic synapses (DDS) are located in the distal dendrites of the GCs, where they also receive dense inhibition from local dSACs (Eyre et al., 2008; Burton and Urban, 2015). In contrast, the proximal dendritic compartment and soma of the GCs receive dense innervation from top-down afferent axons (Záborszky et al., 1986; Matsutani and Yamamoto, 2008; Zaborszky et al., 2012; Nagayama et al., 2014), including excitatory cortical feedback, and cholinergic and GABAergic inputs from the BF (Matsutani and Yamamoto, 2008; Gracia-Llanes et al., 2010; Boyd et al., 2012; Markopoulos et al., 2012; Nunez-Parra et al., 2013; Villar et al., 2021). This synaptic organization of inputs suggests two separate functional domains of inhibition in GCs: a proximal domain, largely containing top-down GABAergic inputs from the BF (Eyre et al., 2008; Villar et al., 2021), and a distal domain, comprised by local feedforward inhibition from dSACs (Eyre et al., 2008). Using a waveform clustering approach, we were able to distinguish between these two populations of inhibitory currents based on their amplitudes and decay times. Our results with selective optogenetic control of proximal (LRGNs) and distal (dSACs) inhibitory inputs onto GCs are consistent with this idea. While LRGNs activation evoked IPSCs with larger amplitude and slower decay time, disynaptic stimulation of dSACs elicited IPSCs with smaller amplitude and faster decay times (clusters 3 and 1, respectively). Segregated inhibitory inputs onto GCs are supported by previous studies indicating that a subpopulation of dSACs preferentially target the distal dendrite of GCs (Eyre et al., 2008), while BF-LRGN axons are notoriously more abundant in the GCL, suggesting a perisomatic innervation of GCs (Gracia-Llanes et al., 2010; Villar et al., 2021). Nevertheless, the amplitude and decay times of the IPSCs evoked by these stimulations overlap to a considerable degree, which comprised cluster 2, suggesting topologically overlapping inputs from LRGNs and dSACs along the GC somatodendritic axis (Burton and Urban, 2015). Intriguingly, the increase in frequency of inhibitory events was observed in half of the population of GCs examined. The reason for this is unknow but we note that the frequency of inhibitory events was not different between responding and non-responding GCs, and disynaptic responses elicited in GCs occurred with high probability. This suggests that there could be differences in the muscarinic response in dSACs, or differences in the efficiency of eliciting disynaptic and/or muscarinic responses in these neurons in the slice. Further experiments should address this possibility.

While signal dampening by electrotonic conduction predicts a decrease in event amplitude for distal events (Armstrong and Gilly, 1992), the faster decay of these events was unexpected. One possible explanation for this discrepancy is that proximal and distal GABAergic synapses onto GCs could have different properties. For example, the composition of postsynaptic GABA_A_ receptors (GABA_A_R) could vary along the somatodendritic axis of GCs, giving rise to synaptic currents with distinctive kinetics. The identity of GABA_A_R subunits expressed in the proximal and distal dendritic compartment of GCs is unknown, however different combinations of GABA_A_R subunits or posttranslational modifications of receptors have been shown to greatly impact the kinetic of the postsynaptic currents (Laurie et al., 1992; Moss et al., 1992; McDonald et al., 1998; Moss and Smart, 2001; Nusser et al., 2013; Nakamura et al., 2015; Nunes and Kuner, 2015). The observed differences in decay time of currents elicited with photo-uncaging are consistent with heterogeneous properties of GABA_A_R along the somatodendritic axis of GCs. Future experiments characterizing single channel properties of GABA_A_Rs along the somatodendritic axis of GCs could clarify the degree of heterogeneity of GABAergic currents in GCs.

ACh regulates network function by altering the excitation-inhibition balance of neural circuits (Pitler and Alger, 1992; Behrends and Bruggencate, 1993; Wanaverbecq et al., 2007), and in the OB, cholinergic modulation has been shown to play a role in odor discrimination and learning (Doty et al., 1998; Linster and Cleland, 2002; Wilson et al., 2004; Hellier et al., 2012; Chapuis and Wilson, 2013; Hanson et al., 2021). Our results provide evidence supporting that the activation of distinct mAChRs results in a differential modulation of intrinsic and extrinsic sources of inhibition onto GCs. Cholinergic modulation of GABAergic inhibition has been previously shown in the hippocampus (Pitler and Alger, 1992; Hasselmo and Schnell, 1994; Wanaverbecq et al., 2007), thalamus (Antal et al., 2010; Ye et al., 2010) and cerebral cortex (Hasselmo and Bower, 1992; Gil et al., 1997; Kimura and Baughman, 1997; Patil and Hasselmo, 1999; Kimura, 2000; Brombas et al., 2014). Intriguingly, while muscarinic presynaptic inhibition of neurotransmitter release has been reported for glutamatergic (Hasselmo and Sarter, 2011), as well as, GABAergic synapses (Behrends and Bruggencate, 1993; Kimura and Baughman, 1997; Patil and Hasselmo, 1999), in the OB, muscarinic presynaptic inhibition occurred in the BF GABAergic input but not in the excitatory cortical feedback. This differential effect at the somatic level would increase the synaptic weight of the excitatory feedback, and somatic disinhibition of GCs, increasing dendrodendritic and lateral inhibition on M/TCs due to a higher likelihood of generating somatic action potentials in GCs, thus reducing spatial activation of M/TCs and possibly favoring odor pattern separation. Interestingly, a previous study (Mazo et al., 2016) showed that regulation of excitation onto GC by presynaptic inhibition can modulate beta oscillations, suggesting that muscarinic modulation of GC inhibition could also affect oscillatory activity in the OB. Further experiments will be necessary to test these possibilities.

## METHODS

### Animals

All experiments were conducted following the guidelines of the IACUC of the University of Maryland, College Park. Electrophysiological experiments were performed on adult C57BL/6 (JAX, stock #664), *Gad2-IRES-Cre* mice (JAX, stock #010802), *Thy1-EYF-ChR2* (JAX, stock #007612) and *ChAT-tau GFP* (generously provided by Dr. Sukumar Vijayaraghavan) female and male mice, ranging in age from one to four months, obtained from breeding pairs housed in our animal facility.

### Stereotaxic injections

Anesthesia was induced with 2% isoflurane at a rate of 1 L/min and adjusted over the course of the surgery. Body temperature was maintained using a heating pad. Carprofen (intraperitoneal, 5 mg/Kg) was used as analgesic and Betadine as antiseptic. During the surgery, the eyes were lubricated using a petrolatum ophthalmic ointment (Paralube). To express ChR2 in LRGNs, *Gad2-Cre* mice received a stereotaxic injection (200 nL) of the AAV5-CAG-Flex-ChR2-tdTomato adenovirus (Catalog #18917, Addgene) into the MCPO, using the following stereotaxic coordinates (in mm): D/V –5.4, M/L ± 1.63, A/P +0.14. Alternatively, to visualize the innervation of BF-LRGNs across the OB layers, the anterograde tracer AAV5-CAG-Flex-eGFP-WPRE (Catalog #51502, Addgene) was injected into the MCPO as detailed above. For anterograde expression of ChR2 in the cortical feedback projections, C57BL/6 mice were injected with the AAV5-CAG-ChR2-mCherry adenovirus (200 nL, Addgene) in the PC, using the following stereotaxic coordinates (in mm): D/V –3.5, M/L ± 2.8 and A/P 1.6. After surgery, animals were left to recover, and electrophysiological recordings or histological experiments were conducted at 3 weeks or later.

### Histology

Mice were transcardially perfused with cold 4% PFA prepared in 0.1 M phosphate buffer saline (PBS) at pH 7.4. Brains were then harvested and post fixed overnight (ON) at 4°C in the same fixative solution. The brain was sliced horizontally in sections of 50-100 μm, nuclei were stained with DAPI (Catalog #D1306, Invitrogen) and mounted in a solution of Mowiol-DABCO. The mowiol mounting media contained 9.6% w/v mowiol (Catalog #475904, Millipore), glycerol 24% w/v, 0.2 M Tris (pH 6.8), 2.5% w/v DABCO (used as antifade reagent, Catalog #D2522, Sigma) and Milli-Q water. BF cholinergic axons were visualized using the *ChAT-tau GFP* mouse line, in which the intrinsic eGFP expression was amplified by immunohistochemistry. Briefly, free floating brain slices of 50-100 μm were blocked with donkey serum (10%, Catalog #S30-M, Millipore) in PBS supplemented with Triton X-100 (0.1% v/v, Catalog #T8787, Millipore, PBS-T) for 1 h at room temperature (RT). Sections were then incubated ON at RT with a rabbit anti-GFP primary antibody (1:500, Catalog #598, MBL) and 2.5% donkey serum in PBS-T. The primary antibody was washed with PBS-T for at least 30 min before incubation with a donkey anti-rabbit antibody coupled to Alexa fluor-594 for 2 h, at RT (1:500, Catalog #A-21207, Invitrogen). Images were acquired using a Leica SP5X confocal microscope and analyzed using ImageJ (NIH) and a custom written MATLAB software (MathWorks).

### Whole-cell recordings

Horizontal OB slices (250 μm) were prepared as before (Villar et al., 2021). Briefly, slices were prepared in oxygenated ice-cold artificial cerebrospinal fluid (ACSF) containing low Ca^2+^ (0.5 mM) and high Mg^2+^ (3 mM). Sections were then transferred to an incubation chamber containing normal ACSF (see below) and left to recover for at least 30 min at 35°C, before the recordings. In all experiments the extracellular solution is ACSF of the following composition (in mM): 125 NaCl, 26 NaHCO_3_, 1.25 NaH_2_PO_4_, 2.5 KCl, 2 CaCl_2_, 1 MgCl_2_, 1 myo-inositol, 0.4 ascorbic acid, 2 Na-pyruvate, and 15 glucose, continuously oxygenated (95% O_2_, 5% CO_2_) to give a pH 7.4. Neurons were visualized with an Olympus BX51W1 microscope using a 40x water immersion objective (LUMPlanFI/IR, Olympus) and recorded using a dual EPC10 amplifier interfaced with the PatchMaster software (HEKA, Harvard Bioscience). Whole-cell recordings were performed at RT. Patch pipettes were made of thick wall borosilicate glass capillaries (Sutter instruments, 3-6 MΩ resistance) using a horizontal pipette puller (P-97, Sutter Instrument). Spontaneous inhibitory postsynaptic currents (sIPSCs) were recorded with pipettes filled with an internal solution of the following composition (in mM): 125 Cs-gluconate, 4 NaCl, 10 HEPES-K, 10 Na phosphocreatine, 2 Na^+^-ATP, 4 Mg^2+^-ATP, and 0.3 GTP. Alternatively, the sIPSCs were recorded at −70 mV using an internal solution of the following composition (in mM): 150 CsCl, 4.6 MgCl_2_, 0.1 CaCl_2_, 10 HEPES, 0.2 EGTA, 4 Na-ATP, 0.4 Na-GTP and 5 QX-314. The final pH of the internal solution was adjusted to 7.3 with CsOH. The measured osmolarity was ∼290 mOsm. For current clamp recordings, the internal solution had the following composition (in mM): 120 K-gluconate, 10 Na-gluconate, 4 KCl, 10 HEPES-K, 10 Na phosphocreatine, 2 Na^+^-ATP, 4 Mg^2+^-ATP, and 0.3 GTP adjusted to pH 7.3 with KOH. For optogenetic stimulation, a LED lamp (COP1-A, Thor Labs) was used to produce brief pulses (0.5-5 ms) of collimated blue light (473 nm, 1 mW/mm^2^) delivered through a 40x water immersion objective.

### Single photon GABA uncaging

We performed single-photon GABA uncaging as previously described (Nunez-Parra, et al. 2013). To visualize the morphology of the recorded neurons we included Alexa fluor-594 (20 μM, Invitrogen) in the recording pipette and used custom written ImageJ μManager software to track and aim the uncaging laser spot along the somatodendritic axis of the GC. The recorded sequence of coordinates was interpolated and recreated by discretized movements of a motorized stage from the proximal to distal coordinates (in ∼ 10 μm steps). Similar results were obtained when the uncaging sequence was reversed to start at the most distal coordinate instead. The collimated output of a 405 nm laser (Coherent, LLC) was expanded to 60% of the back aperture of a 60x Olympus objective. The spot has a Gaussian profile in the focal plane with a 1/e^2^ radius= 0.87 μm. Fluorescence illumination was achieved using a green LED (exciter 594 nm center wavelength) (Chroma), and the emitted light was collected by a CCD camera (Hamamatsu). The concentration of DPNI-GABA was 2 mM (Tocris). Laser flashes were of 100 μs duration with power intensities at the surface of the slice up to 2 mW/μm^2^.

### Data analysis

Electrophysiological recordings were analyzed in MATLAB (MathWorks) and Igor Pro (WaveMetrics). Synaptic currents were detected and analyzed with custom written script in MATLAB. Rise time was calculated as the time from 10% to 90% of the current peak. The decay time was measured by fitting a double exponential decay function to the current relaxation and computing the weighted time constant (*τ*_w_) as *τ*_w_= (a_1_*τ*_1_ + a_2_*τ*_2_)/(a_1_ + a_2_), where a and *τ* are the amplitude and time constant of the first (1) and second (2) exponentials, respectively. The IPSC waveforms from all cells were used to cluster the GABAergic events using the k-means clustering function in MATLAB. To determine the number of clusters we calculated the average distance from each point to every centroid (*d*-value), while varying the number of clusters from 1 to 10. The *d*-value was plotted against the number of clusters used for clustering, and the cutoff for the number of clusters was obtained when *d*-value plateaued (Otazu et al., 2015) (**Figure 2-1B**).

Data is shown as the mean ± S.E.M, unless otherwise specified. Statistical significance was determined by student’s t-test or Wilcoxon rank sum (*= p<0.05, **= p<0.01, ***= p<0.001).

### Pharmacological agents

Drugs were prepared from stocks stored at –20°C, diluted into ACSF and perfused at a speed of ∼2 mL/min; acetylcholine chloride (Catalog # A6625, Sigma), muscarine iodide (Catalog #3074, Tocris), oxotremorine (Catalog #1067, Tocris) 4-DAMP (Catalog #0482, Tocris), AFDX-384 (Catalog #1345, Tocris), pirenzepine dihydrochloride (Catalog #1071, Tocris), atropine (Catalog #A0132, Sigma), nicotine ditartrate (Catalog # 3546, Tocris), PNU-120596 (Catalog #2498, Tocris), 6-cyano-7-nitroquinoxaline-2,3-dione (CNQX disodium salt, Catalog #1045, Tocris), (R)-Baclofen (Catalog #0796, Tocris).

## ACKNOWLEDGMENTS

This research was supported by National Institutes of Health (NIH)-National Institute on Aging Grant AG-049937A (R.C.A) and National Science Foundation-Graduate Research Fellowships Program/Division of Graduate Education Grant 1322106 (R.H). We thank Drs. Rodrigo Andrade and Richard S. Smith for helpful comments on this manuscript and Eric Segev for assistance with the laser uncaging software.

## AUTHOR CONTRIBUTIONS

P.S.V, R.H and R.C.A designed research; P.S.V, R.H and B.T performed research; P.S.V, R.H and B.T analyzed data; P.S.V and R.C.A wrote the paper.

## EXTENDED FIGURES

**Figure 2-1.**
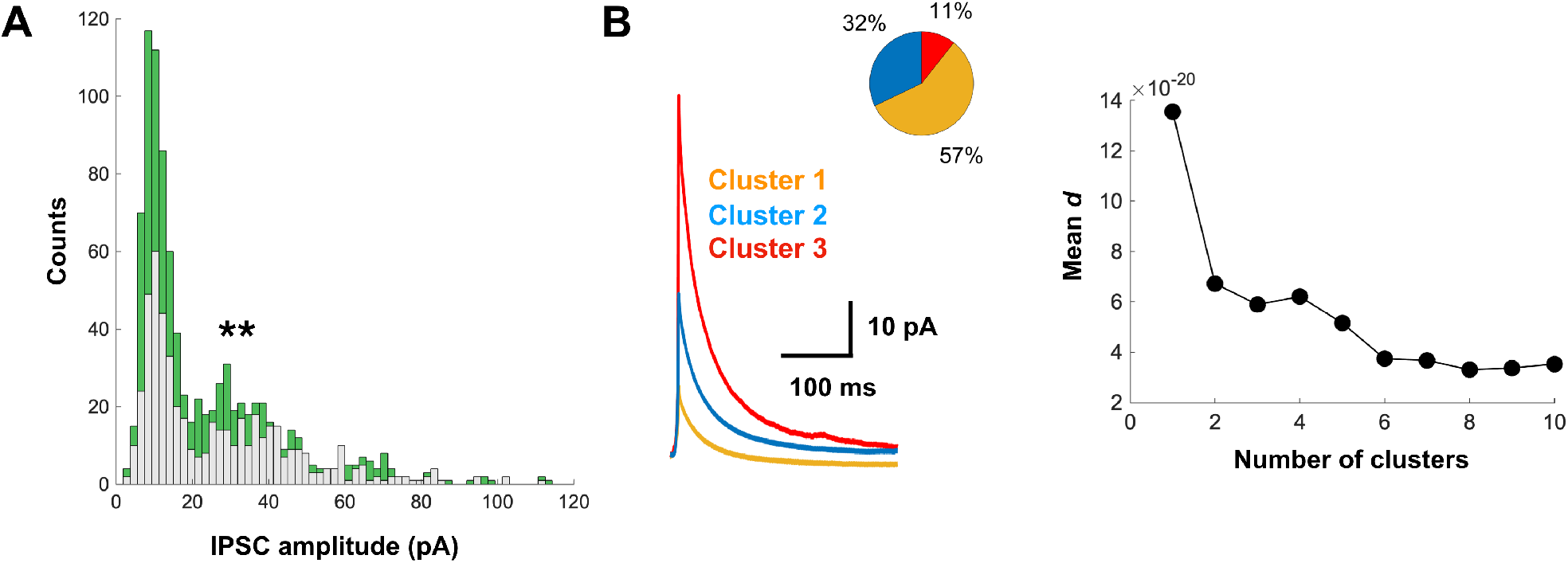
Spontaneous IPSCs in GCs can be sorted into three populations **(A)** Amplitude distributions of sIPSCs, before (light gray) and after Mus application (green, p< 0.01). Mus produces a relatively larger increase in events of smaller amplitude. **(B)** Left, sIPSC waveforms were clustered into three populations using the k-means method. Overlay of the average waveform obtained from the arithmetic mean of all sIPSCs assigned to a particular cluster (cluster 1: n= 808, amplitude 13.9 ± 6.3 pA, rise time 3.2 ± 2.7 ms, decay time 62.5 ± 35.4 ms; cluster 2: n= 453, amplitude 28.6 ± 8.3 pA, rise time 1.7 ± 1.4 ms, decay time 79.5 ± 36.3 ms; cluster 3: n= 150, amplitude 55.2 ± 15.8 pA, rise time 1.3 ± 0.3 ms, decay time 74 ± 29.3). The percentage of events assigned to each of the three clusters is shown in the pie chart. Right, dot plot of the mean *d* value obtained while varying the number of clusters used to determine the number of clusters.

## Notes

Conflict of interest: The authors declare no conflict of interests.

### Competing Interest Statement

The authors have declared no competing interest.

